# Field and mesocosm methods to test biodegradable plastic film under marine conditions

**DOI:** 10.1101/2020.01.31.928606

**Authors:** Christian Lott, Andreas Eich, Boris Unger, Dorothée Makarow, Glauco Battagliarin, Katharina Schlegel, Markus T. Lasut, Miriam Weber

## Abstract

The pollution of the natural environment, especially the world’s oceans, with conventional plastic is of major concern. Biodegradable plastics are an emerging market bringing along potential chances and risks. The fate of these materials in the environment and their possible effects on organisms and ecosystems has rarely been studied systematically and is not well understood. For the marine environment, reliable field test methods and standards for assessing and certifying biodegradation to bridge laboratory respirometric data are lacking. In this work we present newly developed field tests to assess the performance of (biodegradable) plastics under natural marine conditions. These methods were successfully applied and validated in three coastal habitats (eulittoral, benthic and pelagic) and two climate zones (Mediterranean Sea and tropical Southeast Asia). Additionally, a stand-alone mesocosm test system which integrated all three habitats in one technical system at 400-L scale independent from running seawater is presented as a methodological bridge. Films of polyhydroxyalkanoate copolymer (PHA) and low density polyethylene (LD-PE) were used to validate the tests. While LD-PE remained intact, PHA disintegrated to a varying degree depending on the habitat and the climate zone. Together with the existing laboratory standard test methods, the field and mesocosm test systems presented in this work provide a 3-tier testing scheme for the reliable assessment of the biodegradation of (biodegradable) plastic in the marine environment. This toolset of tests can be adapted to other aquatic ecosystems.

## Introduction

Biodegradable plastic materials are being introduced into the market with a growing share in recent years [1], with common uses including food packaging, very lightweight bags and agricultural applications. The possible environmental benefit of biodegradable plastics is dependent on the application. For example, certified soil-biodegradable mulch films [2] are utilized to substitute conventional plastic polyethylene films, especially for those applications in which complete recollection of the film is not possible. After use, certified soil-biodegradable mulch films are plowed into the soil and biodegraded by microbes, avoiding the accumulation of non-degradable plastic fragments in the fields. It is fundamental to keep in mind that biodegradation is not only the visual disappearance or loss of weight of the material, but its remineralization to CO_2_ (and/or CH_4_ under anoxic conditions) and water, and the conversion to biomass by the metabolic action of microbes. Furthermore, biodegradable substitute materials are opted for as a mitigation against marine plastic pollution (e.g. [3]).The proof of environmental biodegradability, i.e. beyond controlled lab test conditions, is crucial for a polymer to be considered as a sustainable alternative to non-biodegradable materials in a specific environment. However, while there exist certification schemes for industrial compostability (e.g., based on standards [4–6]) or soil biodegradability [2], at the time this work was started, there was a lack of standards and methods for assessing marine biodegradability. It is well recognized that positive experimental results on biodegradation of plastics in one environment (e.g. in industrial compost) do not automatically imply comparable or sufficient biodegradation rates of the same material in another system (e.g. in the marine environment) [7, 8].

The knowledge on the biodegradation of biodegradable polymers in freshwater and marine environments is still very limited and experimental data vary strongly. Hence, there is a clear need (and also clear international political call, e.g. The European Green Deal, 12.11.2019 [9]) for reliable environmentally relevant standardized test methodologies to produce comparative and scientifically valid results and for the verification of claims based on sound knowledge rather than assumptions [10]. Data from such tests will help to assess environmental benefits and potential risks and will allow for their comprehensive life cycle assessment.

This need was also emphasized in a recent review [11] which identified three major knowledge gaps regarding biodegradability standards for carrier bags in aquatic environments: the relation of laboratory-based data to patterns of biodegradability within open aquatic environments, the absence of biodegradability standards and test methods for unmanaged aquatic environments and a lack of wider laboratory- and field-based research into polymer degradation within several environments. All three aspects were addressed methodologically in our study.

Biodegradation tests on biodegradable polymers in open aquatic environments have been performed since the onset of industrial development of biodegradable polymers in the 1980s, e.g. of PHA bottles in a Swiss lake [12], films of different starch blends in a river and a pond of Illinois, USA [13], films and plates of different PHAs at the sea shore of Jogashima, Japan [14] and PHB and PHBV test sticks (‘dog bones’) in freshwater ponds, a canal and a seawater harbor in Belgium [15]. Later, marine in-situ tests on several materials in different habitats (surface, open water, sand, mud, mangrove, reef, deep sea) were performed in the Caribbean [16], NW-Atlantic [17], NE Atlantic [18, 19], the Baltic Sea [20–25], NW Pacific [26], NE Pacific [27] SE Asia [28], the Mediterranean Sea [29] and in freshwater [25, 30–33]. Lately, the focus of field research also included microbiological [19, 34] and ecotoxicological aspects [35, 36], and the assessment of the biodegradation of marketed products rather than pure polymers [37–41]. Although field tests have been conducted for several decades the variety of methodologies and experimental conditions render the comparison of results difficult.

*In-situ* experimentation under sometimes harsh marine conditions is challenging and involves several risks including theft, sabotage, conflict with other activities like fisheries and boating, and natural forces such as strong currents and wave action. Loss of samples by anthropogenic impact or natural forces (e.g. [29, 38, 40, 42]) could substantially jeopardize the outcome of experiments. Also, most studies have only tested in one habitat (e.g. [38]) or were too short-termed (e.g. [39]) to produce reliable results on the full biodegradation of a certain material under marine conditions. Many studies also lack the application of a positive control to assess the general microbial activity under the given test conditions and the potential of the microbial community to biodegrade organic materials within the experimentation time at all (e.g. [41]).

The main goal of this study, partly conducted during the EU project Open-Bio [43–45], was to develop robust, reliable *in-situ* test systems to assess the performance of biodegradable plastic materials under natural marine conditions that: (a) withstand natural forces for extended times of exposure, (b) allow for testing in different marine habitats without the loss of samples and (c) provide samples ideally deteriorated by mere biological processes rather than physical destruction. Additionally, the aim was to create a basis for an internationally recognized standard, e.g. EN or ISO, which recently was successfully achieved [47].

When we started this work, standard test methods for *in-situ* testing of plastic materials only existed for the sea surface [46]. For laboratory testing several standards addressing different habitat scenarios are available (water: [48]; sediment-water interface: [49, 50]; beach sand: [51, 52]) and more are under development for the water column [53, 54]. Mesocosm (or tank) tests under controlled conditions simulate an environment which better approximates nature than is possible in small-scale laboratory tests. Thus, mesocosm tests can play an important role as a methodological bridge between field and lab tests (3-tier approach), allowing better understanding of the fundamental parameters controlling biodegradation. Mesocosms of various sizes have been used for the assessment of the performance and environmental effects of conventional and biodegradable plastic under aquatic conditions, either in flow-through systems [e.g. 55, 17, 56 – 62, 83] or closed-circuit tanks [e.g. 63 - 67, 25, 68 – 70, 84] with a large variety of technical properties and environmental conditions. Currently, only one standard test exists [71] which describes the degradation of plastic materials in mesocosm tests; however it requires the supply of running seawater. For this reason, we developed a stand-alone mesocosm system independent of the direct access to seawater, combining the simulation of three coastal marine habitats in a compact unit.

The marine environment is extremely complex and different compartments cover a large variety of conditions. Therefore, a set of tests is required to address this variety. Considering that 80% of plastic pollution originates from land [72], we focused mainly on the coastal regions. We chose three scenarios that cover different ecological aspects: the eulittoral, where plastic is washed to the beach and eventually buried in the sand, the pelagic, where plastic is floating neutrally buoyant in the water and the benthic, where plastic is sunken to the seafloor. Additionally, two different locations were investigated, where tests are technically and economically feasible and environmentally relevant. The field test systems were first developed in the Mediterranean Sea (at the islands of Elba and Pianosa, Italy) and further refined in Southeast (SE) Asia (Pulau Bangka, NE Sulawesi, Indonesia).

For biodegradable plastics, biodegradation is typically assessed with respirometric methods in closed lab test systems, where oxygen consumption or CO_2_ evolution (and/or CH_4_ under anoxic conditions) is measured over time (e.g. [48]). Disintegration occurs as a consequence of biodegradation, however fragmentation and disintegration of plastic, especially under field conditions, can also have purely physical (e.g. UV light, heat, abrasion, mechanical stress) e.g. [68] and chemical (hydrolysis, oxidation) causes.

In this study we used the degree of disintegration of the test material as a proxy for the biodegradation under the condition that (a) the material has been proven to biodegrade under laboratory conditions, hence is biodegradable and (b) under the assumption that our test systems are able to exclude physical forces efficiently enough that disintegration is primarily caused by the metabolic action of microorganisms.

## Materials and methods

### Test materials

To validate the newly developed test systems, the disintegration of a polyhydroxyalkanoate copolymer (PHA, MIREL^®^ P5001, Metabolix, Cambridge, USA)

- a bio-based, bacteria-derived, biodegradable material with a film thickness of 85 µm
- was used as a positive control. A non-additivated low-density polyethylene (LDPE, LUPOLEN 2420K, LyondellBasell, USA) film with a thickness of 30 µm was used as a negative control. PE is the most common and most abundant material for light packaging and shopping bags and without prior physical (e.g. UV, heat) treatment is regarded as non-biodegradable [73, also e.g. 74].

### Test systems and settings

#### HYDRA test frame

The materials were mounted in a holder termed the ‘HYDRA^®^ test frame’ (Fig 1) to protect the samples during exposure against physical stress and to prevent loss of larger fragments after the onset of eventual disintegration. In the Mediterranean Sea tests, the film specimens were covered on both sides with a plastic grid of LDPE (POLY-NET, NSW-General Cable, Nordenham, Germany), with diamond-shaped meshes of 4 x 4 mm size. To be in accordance with standard methods for compost [4], after the successful experiments in the Mediterranean Sea a polyester mesh (PES, SEFAR, Heiden, Switzerland) of 2 x 2 mm mesh size was chosen for the next generation of HYDRA test frames and their subsequent application in another climate zone (tropical SE Asia). This also had the advantage of a better retention of smaller fragments. The exposed area of test material not covered by the filaments of the mesh remained almost the same (51% vs. 52%) and the mesh was not tightly adhering to the film to ensure the complete exposure to water. The specimens covered in mesh were fixed between two plastic frames (PE-HD 300, Technoplast, Lahnstein, Germany) with 3 mm thickness and of 260 x 200 mm external and 200 x 160 mm internal dimensions leaving a surface of 320 cm^2^ of material directly exposed. The frames, the mesh and the test material were then assembled with plastic bolts and nuts (Nylon 6.6, www.kunststoffschraube.de, Singen, Germany) and each test frame was individually tagged with a code made from cable markers (HellermannTyton, Tornesch, Germany) fixed with a cable tie. The dimensions of the test items were chosen to provide enough material for subsampling (Fig 1, left) for further analyses.

**Fig 1.**
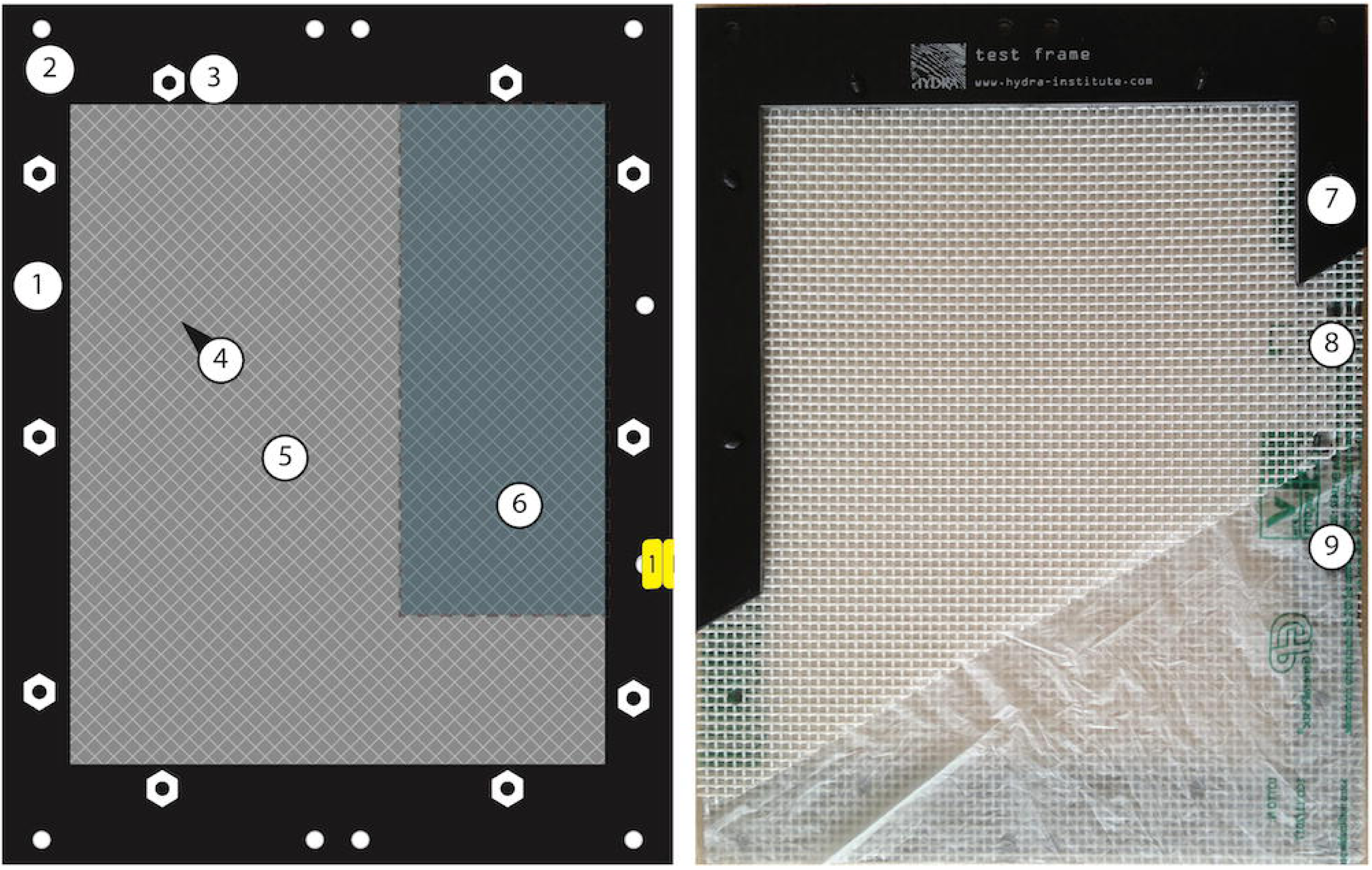
HYDRA^®^ test frame. Left panel: Scheme of test frame. (1) upper frame, (2) holes for fixing the frame to sample holders, (3) plastic nuts, (4) mesh covering the test material, (5) test material (film), (6) area for subsamples. Right panel: photo of HYDRA test frame, cut open to show different layers. (7) upper frame, (8) upper protective polyester mesh (modified to 2 x 2 mm), (9) test material (plastic bag), seen through film: lower mesh, lower frame.

#### Field tests

Three habitats (eulittoral, pelagic and benthic, Fig 2) were chosen to test the disintegration of biodegradable polymers in the sea, reflecting three common scenarios where (conventional) plastic litter is found in the ocean [72, 75].

**Fig 2.**
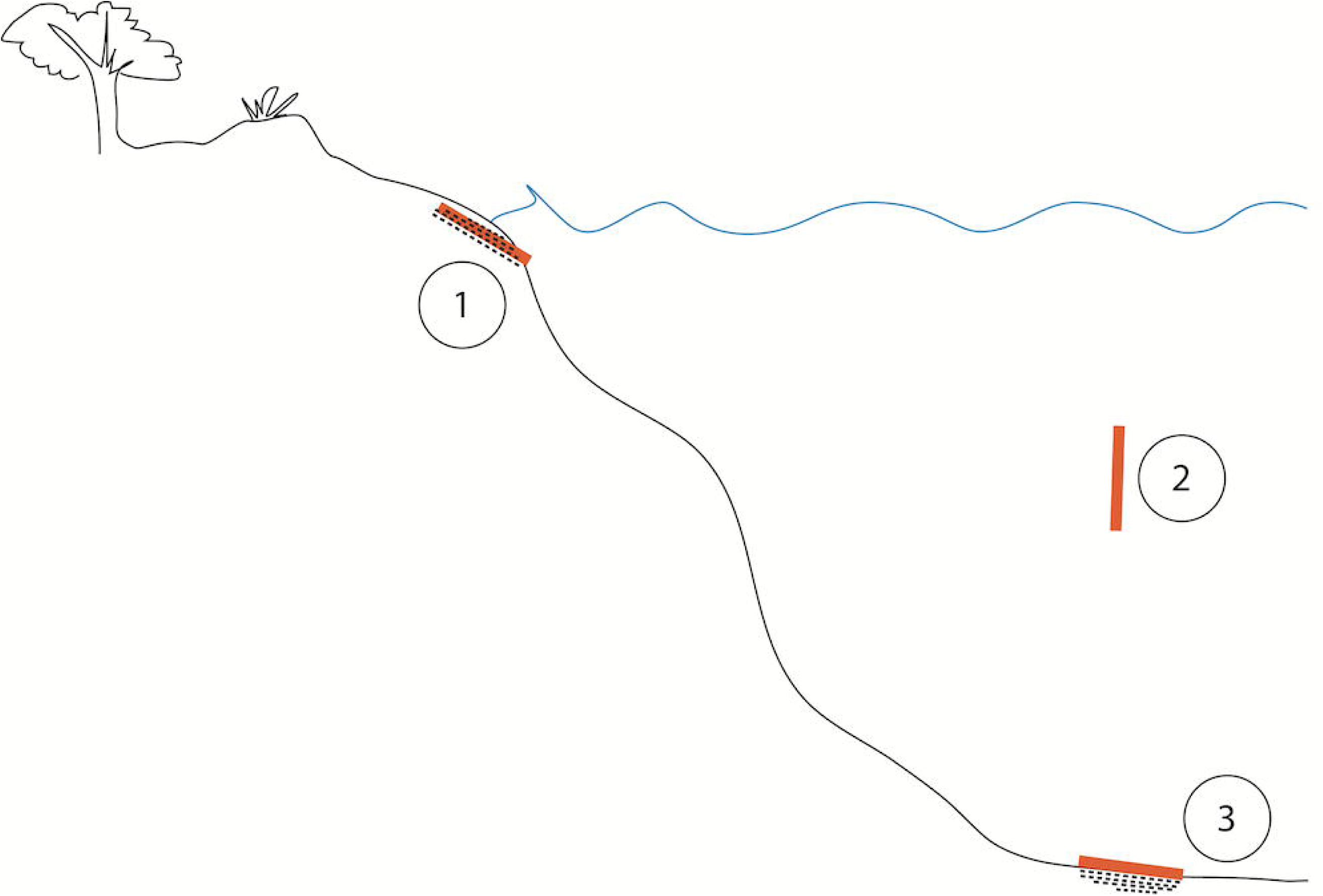
Schematic illustration of coastal habitats. (1) Eulittoral: intertidal beach scenario. Dots around samples symbolize sediment. (2) Pelagic: water column scenario. (3) Benthic: seafloor scenario. Dots below samples symbolize sediment.

For each habitat, a specific support structure for the test frames was developed (see below).

#### Field test locations

##### Matrices: Seawater and sediment as incubation media for the eulittoral (field and mesocosm) and benthic (mesocosm) tests

Natural Mediterranean seawater was taken at the boat slip at Seccheto, Isola d’Elba, Italy (42°44’06.5”N 010°10’33.5”E) and was used to wash the sediment and to fill the mesocosms. Natural marine sediment of siliclastic origin for the eulittoral (field and mesocosm) tests was retrieved at about 0.1 m water depth from the beach of Fetovaia, Isola d’Elba, Italy, (42°44’00.1”N 010°09’15.3”E) and is called “beach sediment”. Carbonate sediment for the benthic mesocosm tests was collected from the seafloor at 40 m depth off Isola di Pianosa, National Park Tuscan Archipelago, Italy, (42°34’41.4”N 010°06’30.6”E) and is called “seafloor sediment”. The sediment was wet sieved with seawater through a 10 mm mesh in order to eliminate coarse particles such as stones and shells and was resuspended several times to flush out very fine particles.

### Study 1: Field experiments Mediterranean Sea

From April 2014 to October 2016 the eulittoral test system as described below was set up in a former saline basin on the Island of Elba, Terme di San Giovanni, Portoferraio (42°48’12.1”N 010°19’01.0”E). This location was chosen because of its unique structure: it is protected against waves and storms by a seawall while still interacting with the open sea by means of several inlets. There is a small irrigation trench that occasionally brings some freshwater runoff to the basin. The ecological character of the site is that of a typical Mediterranean coastal lagoon with weak estuarine influence. Beach sediment was prepared for the eulittoral tests as described above.

The pelagic and benthic tests were performed from June 2014 to October 2016 in the marine protected area of the National Park Tuscan Archipelago off the island of Pianosa (42°34’41.4”N 010°06’30.6”E). The ecological character of this site is that of a typical Mediterranean coastal habitat with low anthropogenic impact and the risk of loss or disturbance by human activities was regarded as very low.

#### Eulittoral field test system

This test system mimicked an intertidal sandy beach in which plastic was buried and experienced changing conditions of being wetted and falling dry with the fluctuations of the sea level caused by tides and waves (Fig 3). A plastic (PP) bin of approx. 60 L volume was used to confine the sediment and the samples. The bottom of the bin was perforated with holes of 25 mm diameter to allow the ambient water to infiltrate. Two layers of plastic mesh (PVC-covered glass fiber 1 x 1 mm and polyester gauze 280 x 280 μm were placed above the holes to prevent the loss of sediment. A layer of 17 cm of beach sand was filled into the bin and a plastic pipe of 32 mm diameter, covered by fine mesh at the top opening, was installed as a short-cut for the rising and falling water to facilitate the drainage of the sediment. Three test frames were buried at a sediment depth of 10 cm with an inclination of about 11° from horizontal to allow the overlaying water to run off after each flood event. The filled bins were placed in groups of five on a wooden rack in order to position the test specimens in the mid-water line in a coastal lagoon. The rack structures and bins were covered by wire barbs to prevent disturbance from birds.

**Fig 3.**
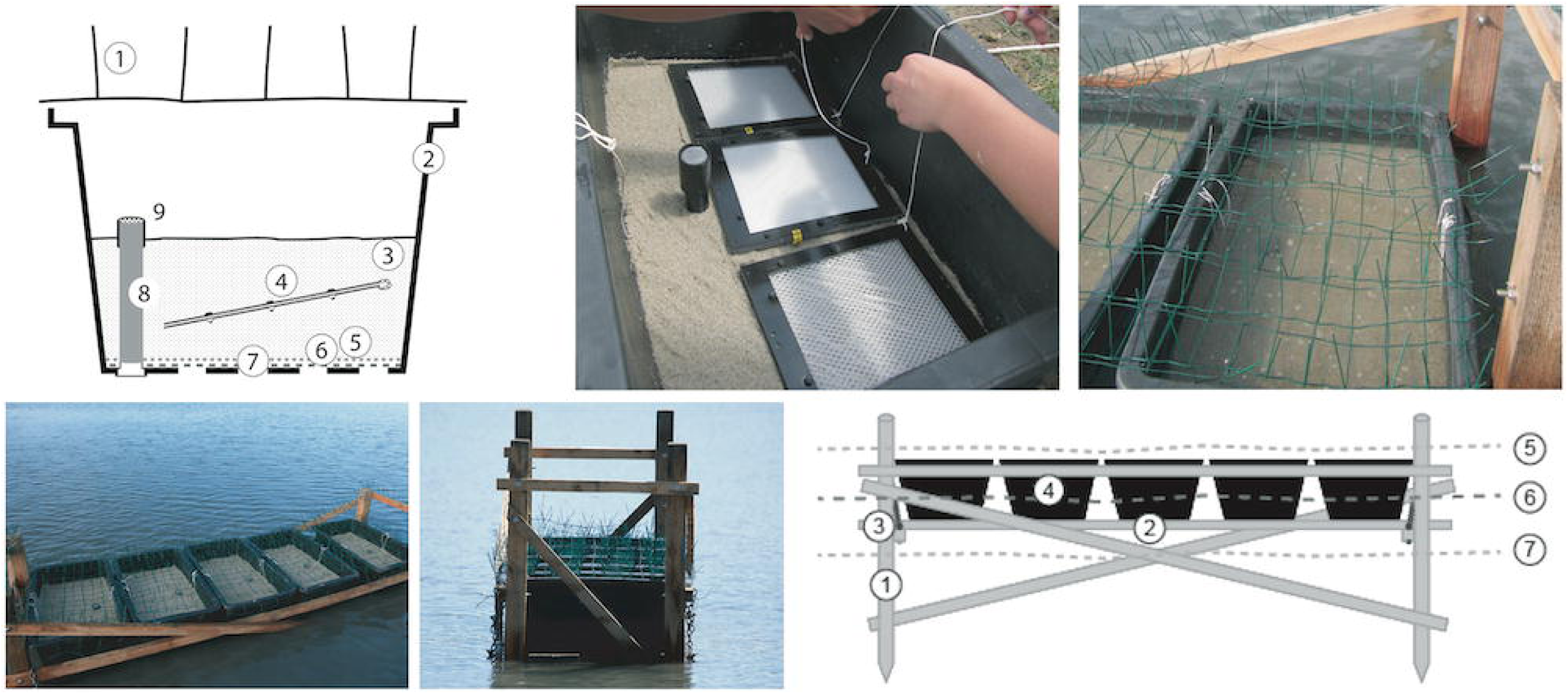
Eulittoral test system. Top left: cross section through bin containing beach sand and samples. (1) wire barbs to deter birds, (2) 60-L plastic bin, (3) sand from a local beach, (4) test item frame, (5) fine mesh (280 µm), (6) coarse mesh (1000 µm), (7) perforations in bottom of the bin, (8) equilibration pipe, (9) fine mesh (280 µm) on the top of the pipe to allow for seawater outflow at high tide. Top center: 3 specimens in the test bin before covering with an additional layer of sand. Top right: test bin at high tide. Bottom left: wooden rack with 5 test bins at midwater. Bottom center: wooden rack with test bins at lower tide. Bottom right: schematic illustration of the eulittoral test system. (1) wooden posts, (2) wooden plank to support the bins, (3) metal chain to adjust the plank, (4) test bin, (5) high water level, (6) mid-water line, (7) low water level.

#### Pelagic field test system

This test system represented the habitat of the free water column where the plastic was neutrally buoyant and was in contact with only seawater as a matrix (Fig 4). The test frames were fixed with cable ties perpendicularly on top of each other to a plastic bar, four of which were mounted to a plastic cylinder (PE-HD 300, Technoplast, Lahnstein, Germany) with stainless steel bolts and nuts to form 4 radial wings. The cylinder was suspended upright at a water depth of about 20 m from a float with a lift of approximately 5 L and kept in place by a concrete anchor weight (50 kg) at the seafloor; this anchor was connected by a stainless steel cable with stainless steel bolts and nuts, or shackles. To prevent corrosion the steel cables were equipped with zinc anodes, fixed with stainless steel bolts and nuts.

**Fig 4.**
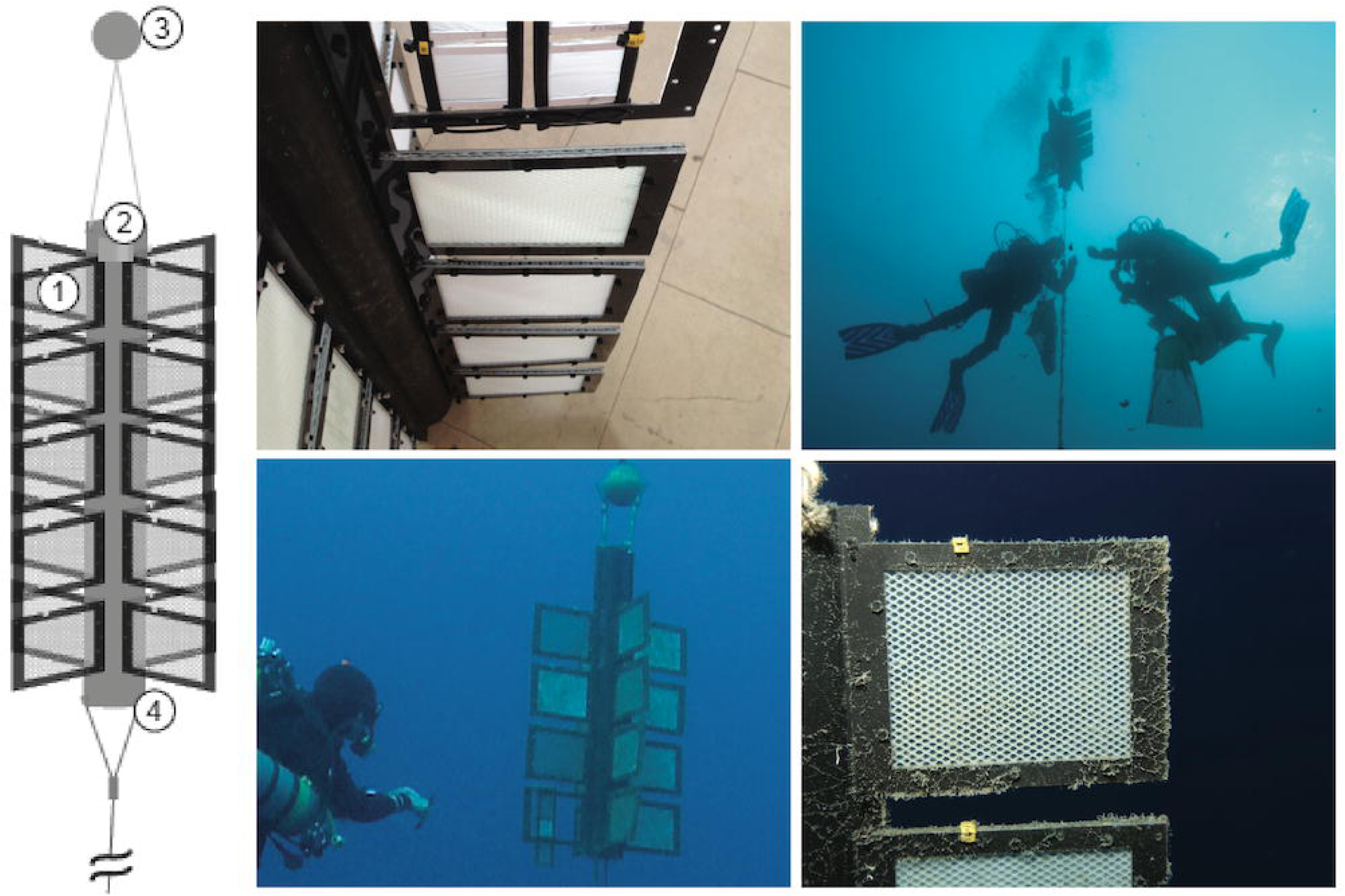
Pelagic test system. Left: schematic illustration. (1) test materials in frames with mesh, (2) plastic pipe suspended from a float at 20 m water depth with specimens radially attached, (3) float, (4) steel cable to anchor stone. Upper left: detail of cylindrical rack with specimens attached radially, upper right and lower left: divers at pelagic test system at 20 m water depth. Lower right: detail of specimen in situ, note biofilm after 2 weeks of exposure (Mediterranean Sea).

#### Benthic field test system

This test system represented the habitat of the seafloor where the plastic was in contact with seawater and sediment as matrices (Fig 5). 3 x 5 test specimens were mounted on a rack of plastic bars and placed on the seafloor at approximately 40 m (Mediterranean Sea) or 32 m (SE Asia) water depth. Each rack was tied to the anchor block of the pelagic test system with a stainless steel cable and fixed to the sediment surface with U-shaped iron bars.

**Fig 5.**
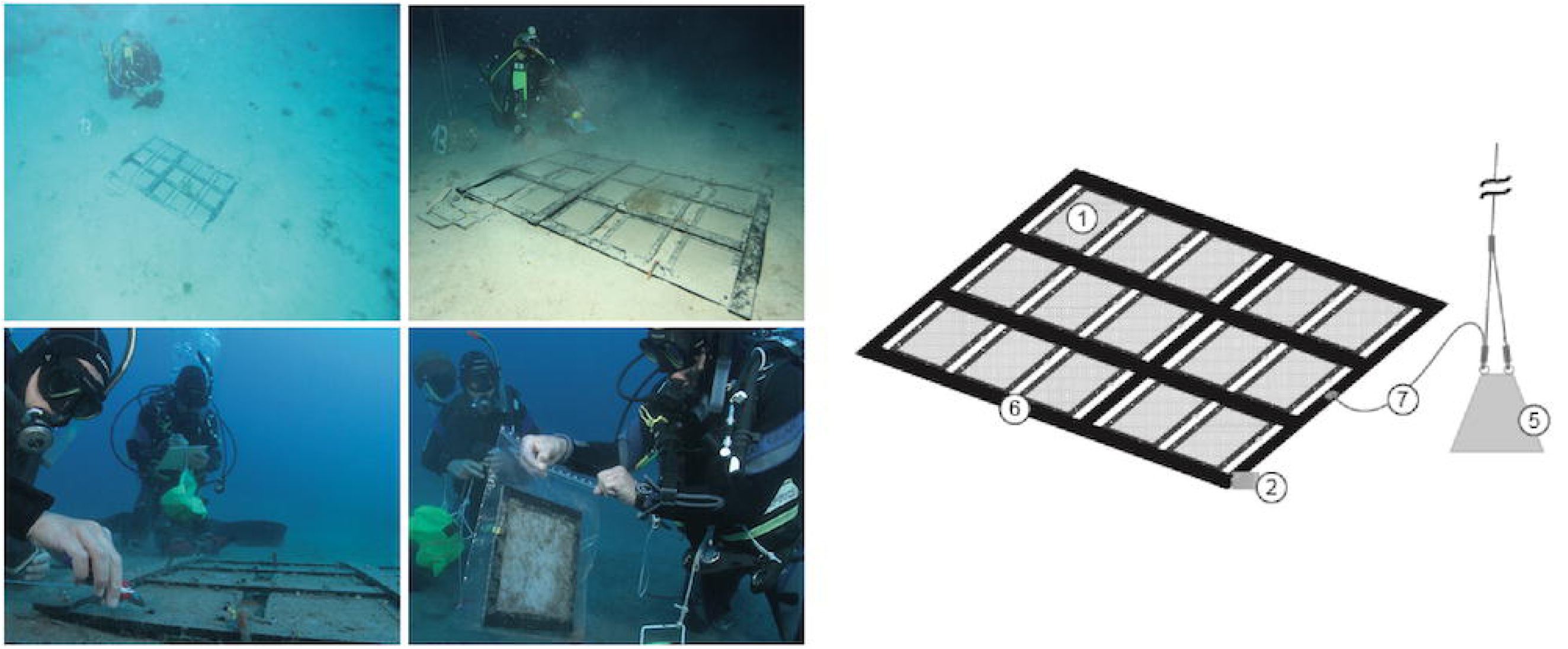
Benthic test system. Right: schematic illustration. (1) test materials in frames with mesh, (2) automated temperature logger attached to rack, (5) anchor stone, also for pelagic test system (see fig 4), (6) plastic rack with specimens in test frames attached, (7) steel cable to anchor stone. Upper row: divers at benthic test system at 40 m water depth (Mediterranean Sea), note sediment and debris on specimens. Lower left: divers detach test frame, fixed with cable ties to the rack. Lower right: specimens are packed individually into PE plastic bags for further treatment in the laboratory.

### Study 2: Field experiments Indonesia

From March 2017 to October 2018 pelagic and benthic tests were performed in Sahaong Bay, Pulau Bangka, NE Sulawesi, Indonesia (01°44’35.4”N 125°09’09.3”E). This location was chosen because it has been used for generations by local people to permanently anchor floating fishing huts (therefore a historical proof of good storm protection). The bay is very narrow and not suited for fishing methods that might have interfered with the test systems such as purse seining or trawling. The benthic system was deployed at a water depth of 32 m on sand to avoid the coral reef. The pelagic test was suspended in 20 m water depth. The ecological character of the site is that of a typical coastal area in the wet tropics, with high water temperature all year round and elevated nutrient content due to terrestrial run-off.

As a modification to the experiments in the Mediterranean Sea the mesh used for all test frames was 2 x 2 mm (Fig 1). There were no eulittoral tests done.

### Study 3: Mesocosm experiments with Mediterranean Sea matrices

The mesocosm experiments simulated, in a multi-100-liter tank under controlled conditions beyond laboratory flask scale (usually a few 100 mL of volume), the degradation of the polymers PHA and LDPE in the marine environment in three coastal habitats: eulittoral, pelagic and benthic (Fig 6). These habitats were mimicked in a combined tank system as follows: PE-HD plastic tanks (Dolav GmbH, Bad Salzuflen) with inner dimensions 93 x 113 x 60 cm were set up in triplicates in a climate room at 21 °C. Each set consisted of the eulittoral test tank placed on top of the tank with the benthic/pelagic test system, connected by a closed-system circuit of about 400 L seawater. Water was pumped into the upper tank in a way that simulated a semidiurnal tide, creating complete flooding every 12 h and a complete draining 6 h later. Tests were conducted twice for about 10 months each and termed year 1 (yr1) and year (yr2), with 4 sampling time intervals each. Before repeating the test, the matrices water and sediment were renewed.

**Fig 6.**
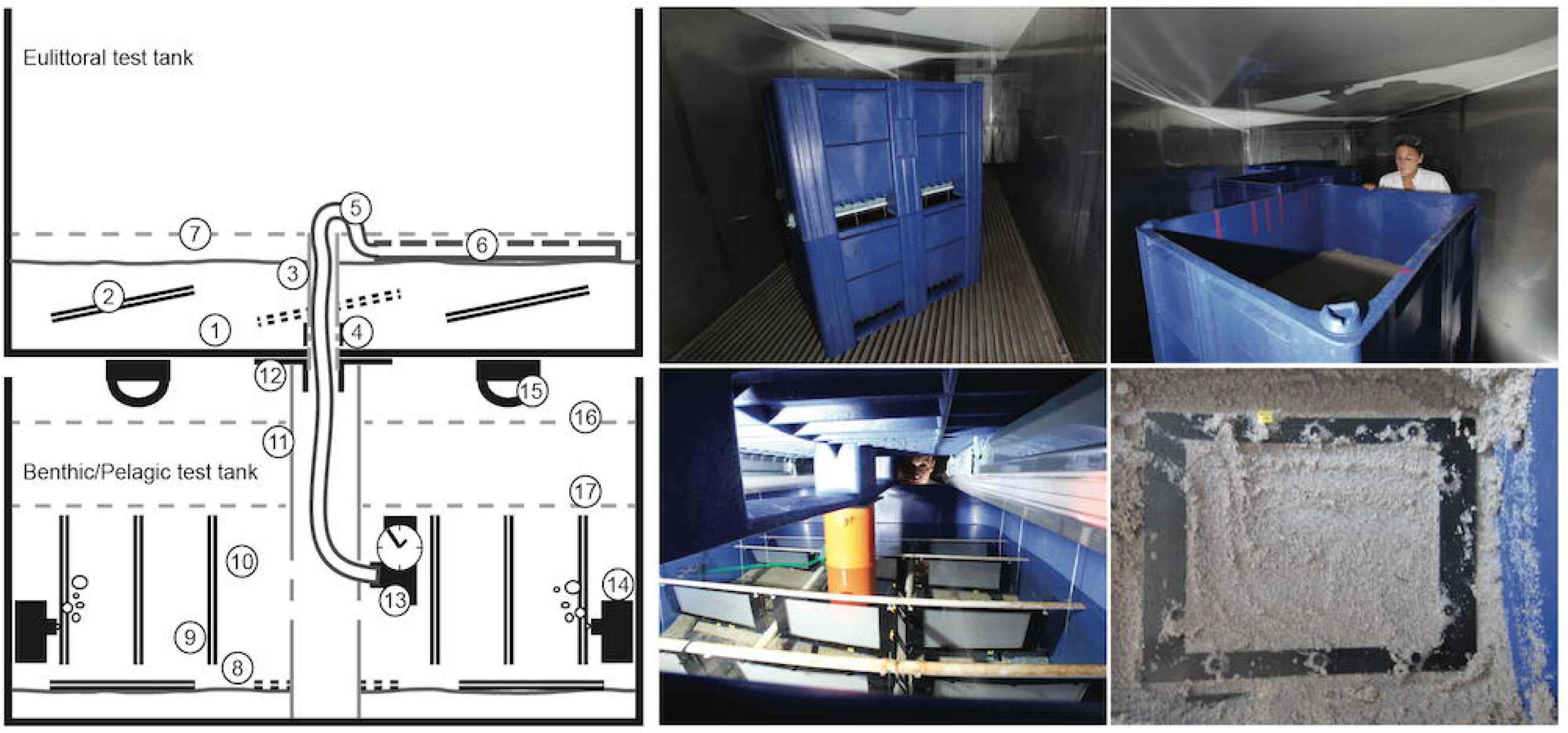
Mesocosm tank system. Left: schematic illustration. Upper left: upper tank, which mimicked the eulittoral (intertidal beach scenario) habitat. (1) beach sand, (2) test frames, (3) drainage pipe connected to lower tank, (4) perforation to drain sediment into (3), (5) hose connected to (13) delivering water to upper tank via (6) a perforated pipe, (7) high-water line in upper tank. Lower left: lower tank which mimicked the pelagic (water column scenario) and the benthic (sublittoral, seafloor scenario, sediment-water interface) habitat. (8) sediment surface, (9) pelagic test frames pending in the water, (10) water, (11) support pipe, connecting lower and upper tank, (12) flange as connector and seal between upper and lower tank, (13) pump with timer delivering water from lower to upper tank, (14) circulation pump for lower tank, (15) aquarium lamps 12h/12h, (16) high-water level, (17) low-water level. Upper center: overview of the mesocosm tank system. Upper right: view into the upper tank with eulittoral test. Lower center: view into the lower tank with pelagic and benthic tests. The pelagic samples were hanging in the water and the benthic samples were placed on the sediment on the tank floor. Lower right: view onto a sample, which was buried in the eulittoral sediment.

#### Eulittoral mesocosm test system

The top tank (Fig 6) mimicked the eulittoral (intertidal) scenario and was filled with a layer of 15 cm of beach sediment. The incoming water was led by a hose to a perforated pipe (diam. 12 mm, L = 1 m) which was laid onto the sediment surface. The water inlet was connected to the benthic/pelagic test tank below by an EHEIM compact 600 pump (Deizisau, Germany) with an adjustable flow of 250 – 600 L/h.

The upper tank had an outlet in the center of the bottom (diam. 50 mm) which was covered by gauze (polyester, 280 µm mesh, SEFAR, Heiden, Switzerland) to prevent sediment loss. The pumping rate was adjusted to balance the inflow with the outflow by passive drainage through the bottom outlet. Twice a day the water level was risen above sediment level by pumping up seawater from the benthic/pelagic test tank below. When the pump was stopped the water was allowed to passively drain through the bottom of the tank, with the consequence of sediment and samples slowly falling dry. Samples mounted in HYDRA^®^ test frames were buried in the sand at 5-10 cm depth with an inclination of 11° from horizontal to prevent water from being trapped on the film at falling water level during the simulated tidal cycles. At the end of a test interval samples were carefully dug out of the sediment and processed as described below.

#### Combined benthic and pelagic mesocosm test system

The benthic/pelagic test tank below the eulittoral test tank (Fig 6) was filled with a layer of 5 cm of seafloor sediment. Test frames were placed horizontally on the sediment surface and weighted down with small blocks of granite to prevent them from being moved by the water flow. The tank was filled with 400 L of natural seawater.

For the pelagic tests, samples mounted in HYDRA^®^ test frames were hung from a rack perpendicularly in the water column. The water was continuously moved by a water EHEIM compact 300 pump with a rate of about 300 L/h. The tank was illuminated from above in a 12/12 h light/dark rhythm by 2 BIOLUX L 36W/965 fluorescent lamps (OSRAM, Munich, Germany) with a nominal luminous flux of 2300 lm each. The water bulk was connected to the eulittoral test set on top and cycled by a pump as described above. At the end of each test interval polymer samples were retrieved from the tanks and processed as described below.

#### Environmental conditions in the different test settings

In order to characterize the natural and the mesocosm habitats the environmental parameters such as temperature, light, salinity, pH, oxygenation were monitored. Additionally, samples of the matrices water, sediment and porewater were taken and analyzed for physical, chemical and biogeochemical properties with standard methods (e.g. [76]). Selected data are given where the environmental conditions are relevant to better understand the methods and their evaluation. The full characterization of the experimental sites will be published elsewhere.

A selection of nutrient-related parameters (C, N, P, Si compounds), photopigments and metals as well as potentially toxic substances such as arsenic and heavy metals (cadmium, chromium, copper, lead, mercury, nickel, zinc), and a catalogue of anthropogenic persistent organic pollutants (POPs) were analyzed in water and sediment from the tanks and from the field sites by external accredited analytical services (Institute Dr. Nowak GmbH & Co. KG, Ottersberg, Germany and SGS-WLN Analytical Services, Manado, Indonesia) according to standard methods.

The chemical analysis was mainly conducted to exclude extreme conditions such as very high nutrients or toxins like heavy metals or POPs in high concentrations that potentially could have inhibited biodegradation. None of the parameters measured was found in high values or unusual of the surrounding biogeochemical context. The detailed list of parameters, methods and detection limits as well as the detailed results will be published elsewhere.

#### Water temperature

Water temperature was automatically recorded with HOBO UA-002-64 Pendant temperature/light loggers (Onset Computer Corporation, Bourne, MA, USA) or TinyTag Aquatic 2 - TG-4100 data loggers (Gemini Data Loggers Ltd, Chichester, West Sussex, UK), attached directly to the test devices.

#### Light

The light regime at the pelagic/benthic field test systems was assessed exemplarily by measuring a depth profile of the photosynthetically available radiation (PAR) within the sensor range of 400 - 700 nm; this was done by a diver from the water surface to the benthic test systems at the seafloor with a LiCor PAR Sensor LI-192 (LICOR Inc., USA) in an underwater housing on sunny days at midday (Mediterranean Sea) or with a HOBO Pendant temperature/light logger in SE Asia. In the Mediterranean Sea the light intensity at the pelagic test system (20 m depth) was 13 % and at the benthic test system (40 m depth) 3 % of the surface intensity. In SE Asia the light intensity at the pelagic test system (20 m depth) was varying between 1.2 and 2 % and at the benthic test system (32 m depth) it was 0.5 % of the surface intensity. UV light which is important as a physical agent for polymer degradation in above water environments is rapidly attenuated with depth under water [e.g. 77, 78] and negligible at the depths of our experiments (pelagic: 20 m; benthic: 32 m (SE Asia)/40 m (Mediterranean Sea)). Thus, it was not measured here.

#### Salinity, pH and oxygen

Salinity, pH and O_2_ concentrations were measured in samples from each of the three field sites: from the free water around the benthic and pelagic test systems, from the sediment porewater retrieved from the bins of the eulittoral test system at the level of the test material, and from the mesocosm tanks. Measurements were taken using a TetraCon^®^ 925 conductivity sensor, a SenTix^®^ 940 pH sensor SenTix^®^ 940, and a FDO^®^ 925 oxygen sensor attached to a Multi 3420 (WTW, Weilheim).

#### Mesocosm tests

The measured water temperature was 20.5±1 °C (mean±standard deviation) (set: 21 °C) and mean light intensity on the sediment surface of the benthic tests was 11.56 μmol photons·m^-2^ ·s^-1^. Salinity was about 39 with a variation of ±1 for both years (set: 38.5). The pH was stable at 8.1±0.1 and the oxygen concentration was close to air saturation (98±2 %). Water chemistry: Throughout the experiments, water in all tanks and porewater from the eulittoral sand contained low to moderate levels of nutrient-related parameters (TN, NO_2_, NO_3_, NH_4_, TP, TOC, DOC, Chl *a*, Phaeophytin, etc.). Compared to the field data nutrient concentrations were similar or only slightly elevated. Dissolved aluminium was detected at low concentration. None of the toxic substances as heavy metals, organotin compounds and POPs were detected. Sedimentology and sediment chemistry: The beach sand of the eulittoral tests was mainly of siliclastic origin and for its grain size was classified as medium sand [79]. The sediment used for the benthic experiments was composed mainly of carbonate minerals and characterized as fine sand in both years. Accordingly, porosity and permeability were slightly lower for both sands in the second year (Table 1) whereas nutrient contents were similar and low. Metal concentrations were low or below detection limit. None of the known anthropogenic toxins analyzed in the sediments used for the experiments were detected.

**Table 1.**
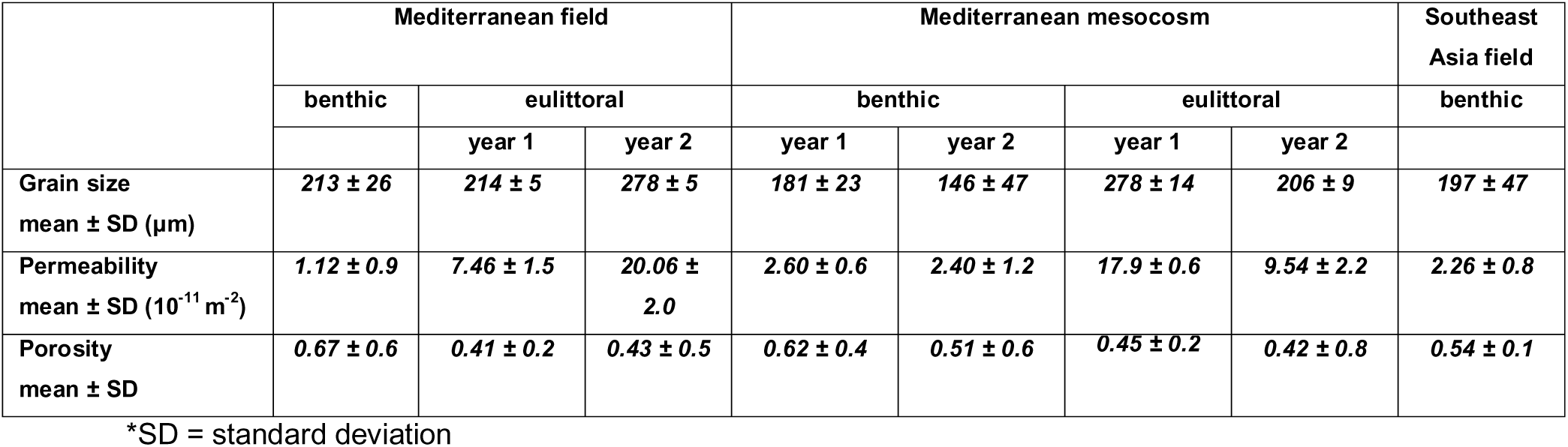
Physical properties of the sediments from the field and mesocosm experiments

#### Mediterranean Sea field test - eulittoral

The temperatures in the eulittoral test bins roughly followed the temperature of the ambient water with the daily and the tidal cycles. Daily mean temperatures of about 25 °C (yr1) and 26 °C (yr2) were highest between June and September and lowest mid-October and March with around 12 °C (yr1 and 2). The lowest daily minimum was −1.0 °C in January 2016. Fluctuations were pronounced, and the night temperatures were usually lower (by up to 5 °C) than in the surrounding water. Salinity of seawater in the test bins was most of the time in the range of 36 – 42; lowest values were almost 0, highest values reached 70. The pH was around 7.6 – 8.2; lowest values were below 6.8, highest values reached 9. The oxygen concentration in the test bins was mostly between 20 and 80 %, and rarely reached 100 % air saturation (see also Table 2).

**Table 2.** Water temperature, salinity, pH and oxygen concentrations of the field and mesocosm experiments.

|  | Mediterranean field |  |  |  | Southeast Asia field |  | Mediterranean mesocosm |
| --- | --- | --- | --- | --- | --- | --- | --- |
|  | eulittoral |  | pelagic | benthic | pelagic | benthic | all habitats |
|  | year 1 | year 2 |  |  |  |  |  |
| Highest daily max. temp. (°C) | 34.5 | 38.4 | 25 | 20 | 28.6±0.4** | 28.6±0.5** | 20.5±1.0** |
| Highest daily mean temp. (°C) | 25 | 30 |  |  |  |  |  |
| Lowest daily mean temp. (°C) | 12 | 11 | 14 |  |  |  |  |
| Lowest daily min. temp. (°C) | -0.8 | -1.0 |  |  |  |  |  |
| Salinity range | 36 - 42* |  | 38 |  | 34.0±0.4** | 33.9±0.2** | 39.0±1.0** |
| Minimum salinity | ~0* |  |  |  |  |  |  |
| Maximum salinity | 70* |  |  |  |  |  |  |
| Mean pH | 7.6 – 8.2* |  | 8.2 |  | 8.1±0.1** |  | 8.1±0.1** |
| Minimum pH | 6.8* |  |  |  |  |  |  |
| Maximum pH | 9.0* |  |  |  |  |  |  |
| Oxygen (% air saturation) | 20 – 80*<br>rarely ~100* |  | ~100 |  | 100.8±4.6** | 101.3±6.8** | 98.0±2.0** |

\*The data for Mediterranean field eulittoral were analyzed from sediment porewater. \*\*mean ± standard deviation.

Water chemistry: The porewater in the bins throughout the experiments had low to moderate levels of nutrient-related parameters, however compared to the benthic and pelagic test sites they were slightly elevated. Dissolved Fe, Mn and Al were detected at low concentrations. None of the POPs tested were detected. Sedimentology and sediment chemistry: The beach sand used in the test bins was fine (mean 214±5 µm yr1) and medium (mean 278±5 µm yr2) sand. The porosity and permeability were therefore higher in the second year (Table 1). Nutrient concentrations were low or b.d.l. and of the metals analyzed only Zn, Ni and Cr were detected in low concentrations. Organotin compounds, pesticides and their metabolites could not be detected in the sediment.

#### Mediterranean Sea field tests - pelagic and benthic

The temperature at the two stations diverged over the course of a year due to stratification of the water column from spring to late autumn, when the temperatures in the different depths merged again. Highest daily mean temperatures were about 25 °C at the pelagic test site and 20°C at the benthic test site. Lowest temperatures were recorded early February until mid-May with around 14 °C at both sites. Fluctuations between daily minimum and maximum temperatures sometimes ranged 4 – 8 °C (pelagic) and 4 – 5 °C (benthic). At the pelagic tests at 20 m depth temperature fluctuations were more pronounced during spring and summer, reflecting an irregular mixing with the surface waters and a gradual downward migration of the thermocline through the water column. At the benthic test, autumn fluctuations were more pronounced due to occasional storms that led to a mixing of the water column. At the depth of the pelagic test system (20 m) solar irradiation was about 13 % and at the benthic test system (40 m) about 3 % of the surface level. Salinity was at 38, pH was 8.2 and oxygen content was around 100 % air saturation at both sites.

Pelagic and benthic water chemistry: The water at both test sites, as well as the porewater at the benthic test site, had low levels of nutrient-related parameters throughout the experiments. Dissolved Al could be repeatedly detected at low concentrations, as well as iron and manganese in the second year. Other metals were b.d.l.. No toxins or POPs were detected. Benthic sedimentology and sediment chemistry: The benthic sand was fine sand (mean 213±26 µm) with low concentrations of nutrients and some metals (Fe, Mn, Pb, Cr). Organotin compounds and pesticides and their metabolites could not be detected. No toxins or POPs were detected in the sediment.

#### SE Asia field tests - pelagic and benthic

The temperature at the two stations was the same throughout the year (pelagic test: 28.6±0.4 °C, benthic test: 28.6±0.5 °C). At the depth of the pelagic test system (20 m) solar irradiation was about 1.5 % and at the benthic test system (31 m) about 0.5% of the surface level. The salinity at the two stations was similar throughout the year. The mean salinity was 34.0±0.4 at the pelagic test, and 33.9±0.2 at the benthic test. A pH of 8.1±0.1 was the same at both test sites, and oxygen concentrations were 100.8±4.6% air saturation at the pelagic test site and 101.3±6.8 % at the benthic test site.

Pelagic and benthic water chemistry: The water at both test sites as well as the porewater at the benthic test site had low levels of most nutrient-related parameters throughout the experiments. The silica, chlorophyll *a* and phaeophytin concentrations however were higher than at the test sites in the Mediterranean Sea. Metal concentrations were all b.d.l., only Cd was detected once in very low concentrations. No toxic substances or POPs were detected. Benthic sedimentology and sediment chemistry: The benthic sand was fine sand (mean 197±47 µm) with low nutrients and low concentrations of some metals (Fe, Mn, Pb, Cr). No toxins or POPs were detected in the sediment.

### Plastic sampling and disintegration analysis

After a given time interval the samples were detached from their racks (pelagic, benthic) or dug out of the sediment (eulittoral), gently rinsed in ambient water, packed individually in PE plastic bags and kept wet until further treatment the same day. In the laboratory, the frame was opened, the material was carefully detached from the upper frame and mesh and photographed immersed in seawater in a tray with a digital camera (Canon EOS 5DMkII, or a SONY A7 III, with a 100 mm or 50 mm macro lens). Subsamples were taken for later optional microscopic and molecular analyses, which are not subject of this publication. The remaining sample was rinsed in deionized water, air-dried and archived.

The disintegration of each specimen was determined photogrammetrically [40]. Dried samples from the Mediterranean Sea experiments were scanned on a flatbed scanner (LIDE 210, Canon Inc.). The samples from the SE Asia experiments were photographed immediately after retrieval. The images thus obtained were analyzed for area loss using ImageJ software (https://imagej.nih.gov/ij/) or GIMP (http://www.gimp.org/) by outlining the boundaries of the remaining test material that had been exposed and not covered by the frame. Subsequently, the ratio of total area vs. lost area was calculated as % area loss.

#### Ethical statement

Sampling of water and sediment in the protected area of the Island of Pianosa was performed with the research permit n.3063/19.05.2014 from the National Park Tuscan Archipelago, Portoferraio. Research in Indonesia was conducted under the research permits no. 71 and 72/SIP/FRP/E5/Dit.KI/III/2017 and extensions granted by the Indonesian Government Ministry of Research, Technology and Higher Education, RISTEK-DIKTI. With regard to the photographs shown for illustration the individuals (all coauthors) in this manuscript have given written informed consent (as outlined in PLOS consent form) to publish these case details.

## Results

### Study 1: Field Mediterranean Sea

All samples were retrieved after the planned exposure time, no sample was damaged nor lost.

#### Visual appearance of the samples

For LDPE, after testing the only visible material alterations were superficial discolorations, presumably from mineral precipitations, biofilm and colored organisms, e.g. sponges. Exemplary images are shown in Fig 7. In the PHA specimens, the first visible signs of disintegration were translucent areas due to progressive thinning of the polymer films, followed by the appearance of cracks, holes and finally fragmentation in all tests. Most samples showed a change in color during the time of the experiment, attributed to the formation of biofilm and mineral precipitates, rendering image analysis difficult in some cases.

**Fig 7.**
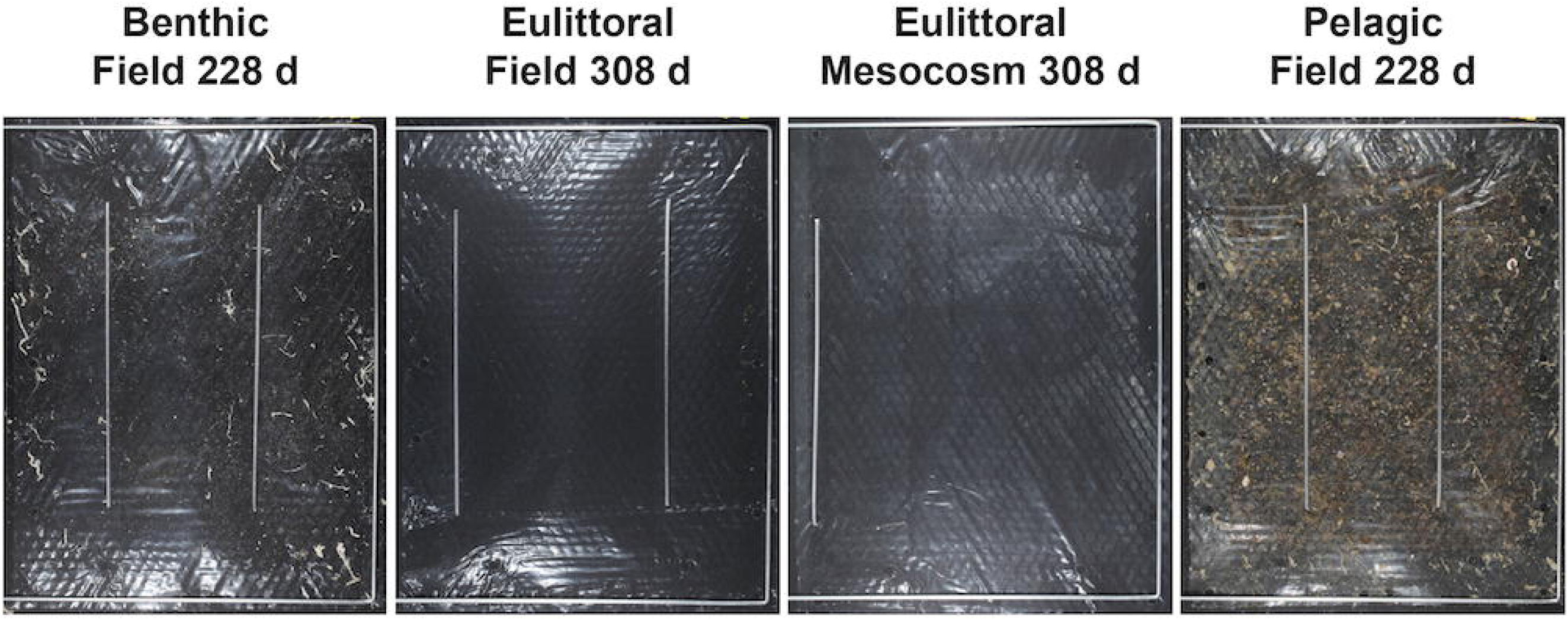
Exemplary photos of LDPE samples from disintegration tests. Freshly retrieved samples from field and mesocosm experiments under Mediterranean Sea conditions. Note: the metal bars in the images were put onto the film to keep the sample under water for photography.

#### Polymer disintegration

For LDPE, disintegration was not detected in any of the tests, and will not be described further in the results section. PHA disintegration was observed in all tests.

#### Eulittoral test (intertidal beach scenario)

PHA disintegrated in both years, reaching more than 50 % after 5 months in yr1 in most replicates. Some replicates however showed only minimal disintegration of between 5 and 20 % after 5 to 10 months. After 22 months all five replicates were disintegrated to over 65 %. In the yr2 experiment PHA disintegration was slower and reached just over 30 % after 10 months for one replicate. All replicates were disintegrated to more than 10 % after 5 months. Generally, the disintegration of all replicates of all sampling intervals was heterogeneous.

#### Benthic test (sublittoral seafloor scenario)

PHA showed disintegration, which differed however by year. One sample showed disintegration of 31 % after 5 months and two samples 60 – 75 % after 22 months. The disintegration of all other samples in both years was below 5 %. In the yr2 experiment, PHA disintegration was slower and only one sample reached more than 6 %. Generally, the disintegration of all replicates was heterogeneous.

#### Pelagic test (water column scenario)

At sampling, all specimens were still intact after the exposure time of max. 22 months. In some specimens, a thinning of the material was observed, visible by more translucent areas. Small holes were detected in only a few samples, but accounted for less than 1 % area loss.

### Study 2: Field tests SE Asia

#### Benthic test (sublittoral seafloor scenario)

In the benthic test of the SE Asia study, PHA showed rapid disintegration, with material loss already well visible after 19 days of exposure (Fig 8). After 90 d most of the test material was gone in all 3 replicates (exemplarily: Fig 8). A conspicuous longitudinal crack was observed in all specimens after exposure, in the cases where there was still material left (Fig 8). This is assumed to have been the consequence of a systematic material irregularity caused during the film processing on a small laboratory-scale extruder rather than on an industrial machine. The crack was not taken into account as loss by disintegration but ignored for the area measurement. The inhomogeneous disintegration of the samples, i.e. higher material loss at the bottom edge might be due to the slight inclination of the seafloor leading to the accumulation of a layer of fine sediment within the test frame building up from the lower part.

**Fig 8.**
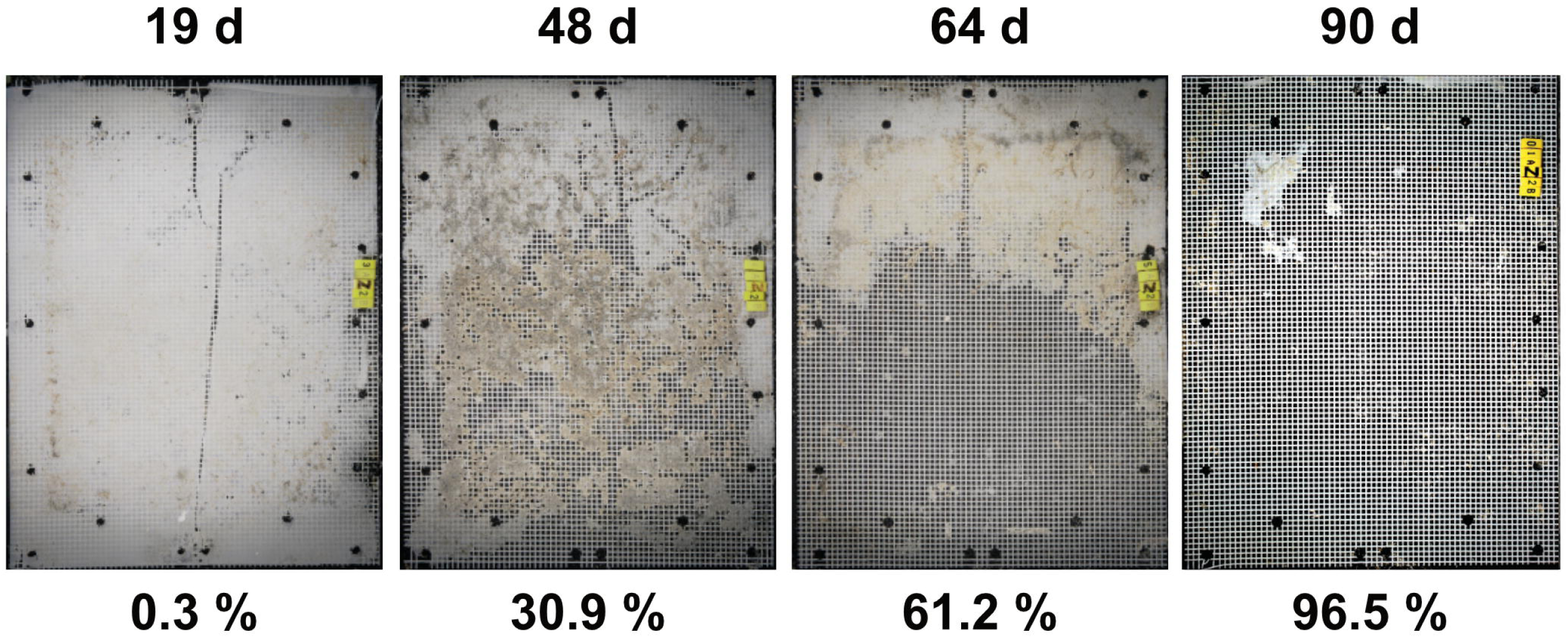
Exemplary photos of PHA samples from benthic field disintegration tests in SE Asia. Photos of 1 replicate from 4 exposure intervals each, with disintegration per sample in % area loss below each image. Note: the values are calculated only from the exposed area not covered by the frame.

#### Pelagic test (water column scenario)

Disintegration was observed in all PHA samples, however, the accurate measurement of the lost area was not possible due to heavy fouling of the support structure and the polymer itself (Fig 9). Also, the layer of the fouling organisms (mainly encrusting red algae, sponges, hydrozoans and bryozoans) often was bonding the mesh and the polymer together. Splitting open the test frame during post-sampling in the lab caused further fragmentation and thus a mechanical destruction of the materials as an artefact. In these cases, both halves of the test frame with the adhering sample were photographed and subjected to analysis (Fig 9). Another factor of sample deterioration in the pelagic tests was linked to animals found in between the meshes. Presumably, crabs and clams had settled as larvae on the test material and after having grown above mesh size they had been trapped between the two layers of protecting mesh. Traces of their movements point to these animals having mechanically destroyed the polymer. Because the degree of error could not be assessed a numerical quantification of the disintegration in the pelagic test in SE Asia is to be treated with caution, and rather seen as an overestimation. It is notable that outside the area of interest, i.e. under the supporting frame where only microbes could biodegrade the samples, almost all material was disintegrated (Fig 9, bottom).

**Fig 9.**
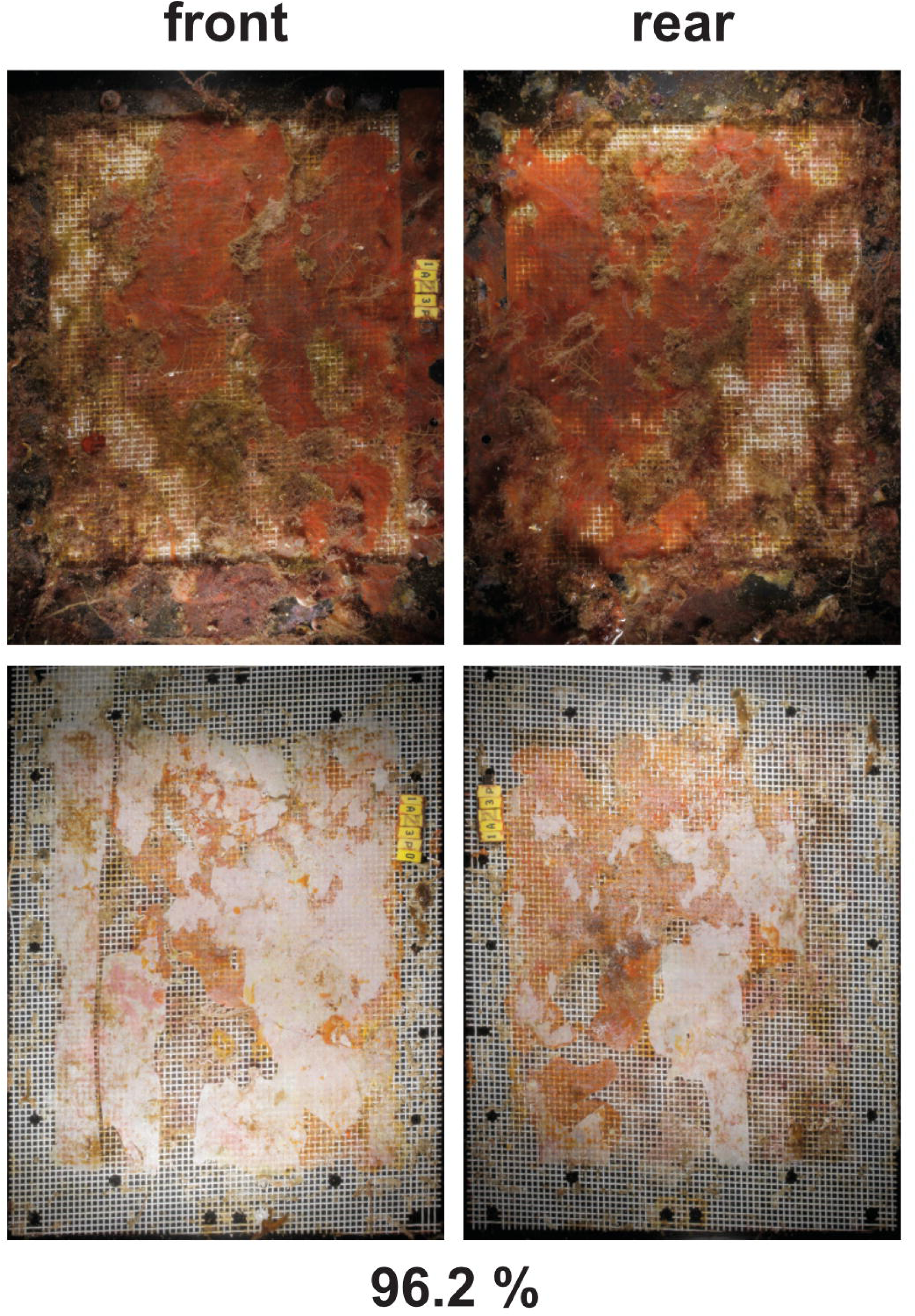
Pelagic field test SE Asia: PHA disintegration. Exemplary photos of a sample after 210 d of exposure. Top: specimens retrieved from the field, note the dense fouling. Images left and right show front and rear of the same sample. B: specimen unpacked, note the almost complete disintegration under the frame, and no fouling there. The light surface is mostly the crust of coralline algae remaining after the disintegration of the test material. The test material is largely disintegrated, as expressed by the percentage of area loss. Note: the value given is rather an overestimation due to the likely mechanical influence of macro-organisms (see main text).

### Study 3: Mesocosm tests with Mediterranean seawater and sediment as matrices

#### Polymer disintegration

PHA samples showed disintegration in all three tested habitats eulittoral, benthic and pelagic. The disintegration gradually progressed with time, with most samples disintegrating less than 50% after 9 – 10 months of exposure. The rate of disintegration differed between habitats and was heterogeneous between replicate samples of one sampling interval, with values ranging e.g. from 7.4 to 46.7 % after 152 d for the benthic test (exemplarily: Fig 10).

**Fig 10.**
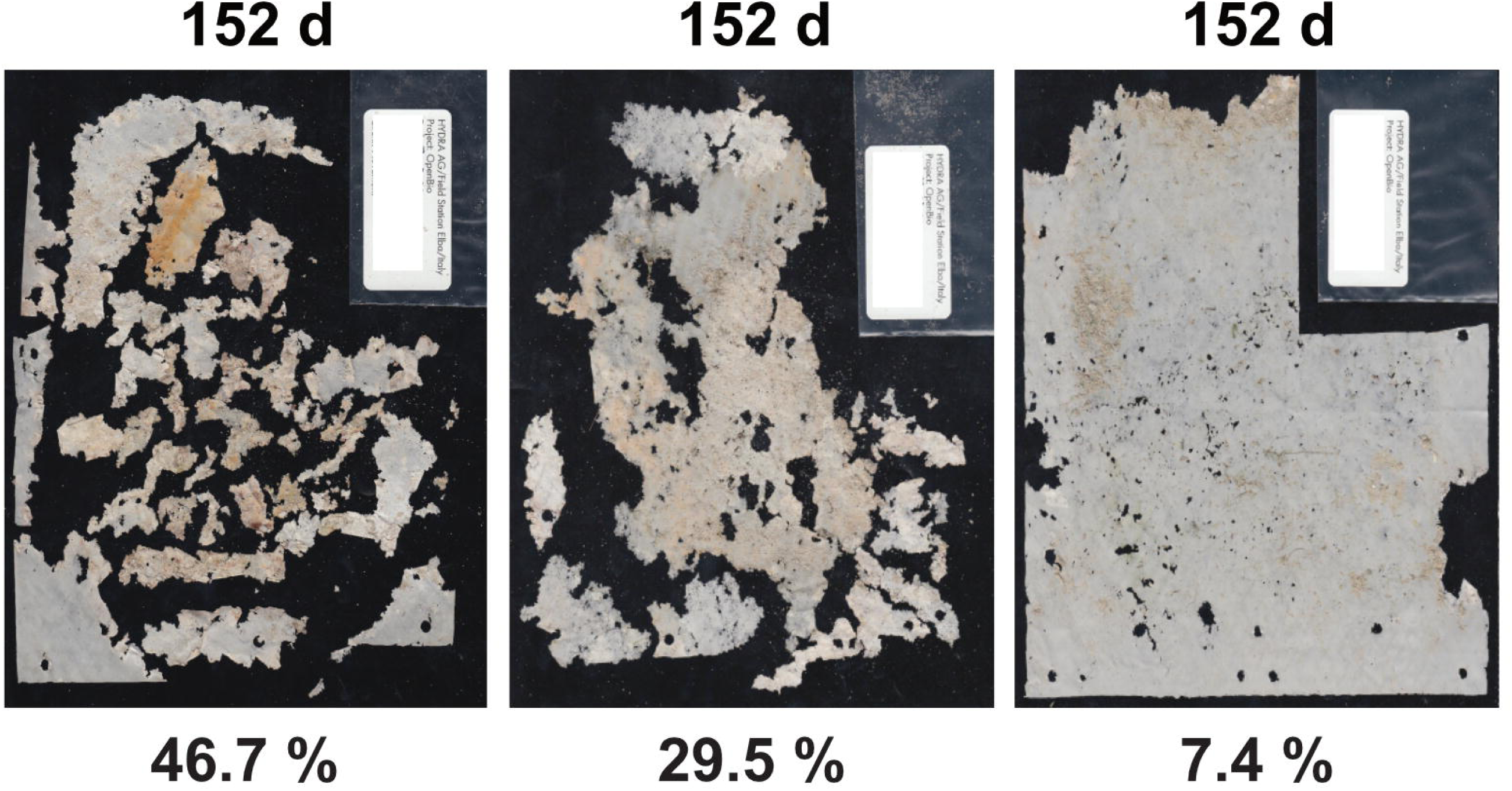
Photos of PHA samples from mesocosm benthic disintegration tests to illustrate the heterogeneity between replicates. Scan images of all 3 replicates from the mesocosm benthic test (sublittoral, seafloor scenario) after 152 d of exposure. Disintegration in % area loss below each image. Note the heterogeneous disintegration between the three replicates.

#### Additional observations

##### Macro-organism interaction with the test materials

Several measures were taken to prevent a negative impact by animals on the test system. However, some disturbances resulted from the interaction of (macro-) organisms with the exposed materials in the field, and to a lower extent also in the mesocosms.

Eulittoral field tests: The picking of birds in the eulittoral test bins was successfully prevented by wire barbs on top of the test system. Infauna like worms or clams were not observed in the eulittoral test bins. A structural problem for the racks made from untreated larch wood resulted from the activity of wood-boring bivalves (“shipworms”) that had destroyed the posts after two years to the brink of collapse (Fig 11).

**Fig 11.**
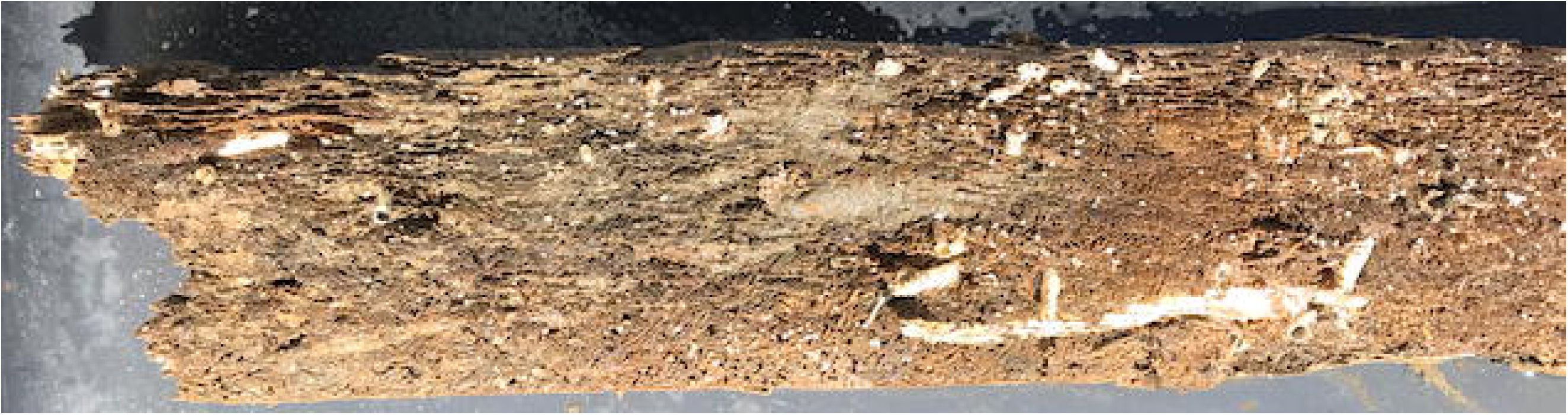
Wooden post from eulittoral test rack. Post from untreated larch wood after 26 months of exposure at the Mediterranean test site. The wood is perforated by wood-boring bivalves (white tubes) and broken. Scale: white tube lower right is about 5 mm diameter.

Pelagic field tests: Fouling, i.e. the colonization of the structures by a community of larger sessile organisms (e.g. macroalgae, sponges, hydrozoa, bryozoan, ascidians, bivalves, tubeworms) was a major issue for the maintenance of the whole structure - from the steel cable to the floats, the plastic support cylinder and the test frames (Fig 9, top and fig 12). Regular cleaning of the support structures (not the samples themselves) by divers was necessary to prevent the test systems from becoming too heavy. At the samples themselves the fouling amalgamated with the test materials, growing through the mesh. For heavily fouled samples it was often impossible to remove the overgrowth after sampling from the remaining polymer without creating further damage and influencing the result. Steel cables were chosen over plastic rope because turtles and some fish are known to snatch ropes while feeding on sponges and other encrusting organisms. However, on several occasions bite marks on the frames and mesh were observed, but a sample was never found bitten all the way to the specimen. On two occasions cable ties that served to fix the codes to the samples were found cut open, and once a large puffer fish was observed grazing on the pelagic test system, having just bitten off a code tag.

**Fig 12.**
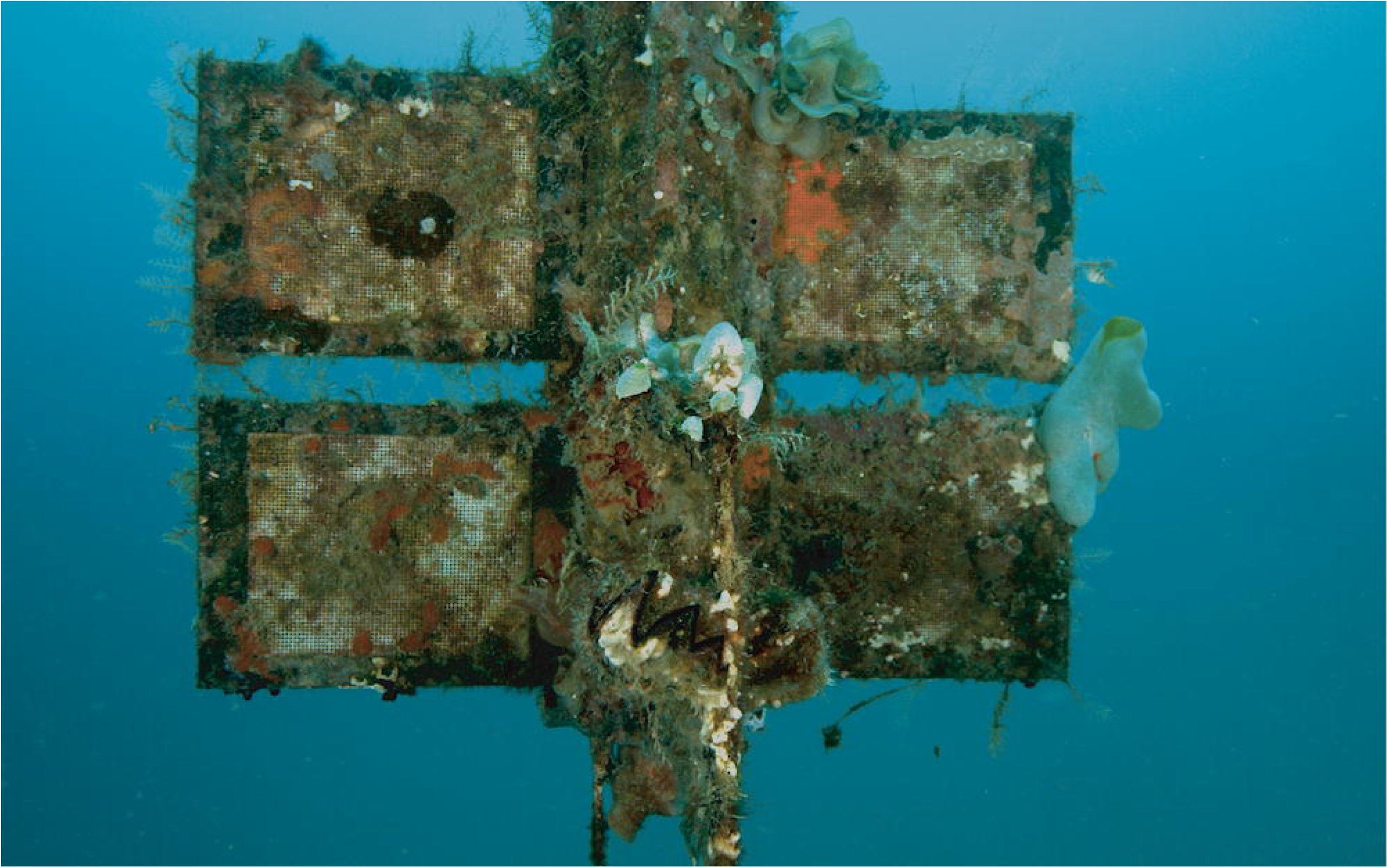
Fouling at pelagic test system SE Asia at −20 m. Sample frames and support structure were covered almost completely by sessile animals and algae. Note the fist-size oyster in lower center (zig-zag line). Exposure time 20 months.

Benthic field tests: The test frames were laid onto the sand bottom and fixed with steel bars. The flat rigid structures were readily populated by benthic fish such as gobies which dug out dens in the underlying sediment. Octopus were observed using the benthic samples as shelter and also sea cucumbers were found digging under the panels. Bioturbation presumably by sand-dwelling worms, bivalves and shrimp under and around the specimens was also observed having changed the sediment topography at the test sites at each sampling event. On some occasions, sediment and/or seagrass leaf debris was found to have accumulated on the benthic samples, but was absent at the next maintenance dive, indicating some water dynamics at the water-sediment interface.

## Discussion

### Field test systems allow for long-term observation under natural conditions

Three scenarios were chosen to mimic conditions at locations where dispersed plastic debris is commonly found in coastal marine areas: the beach scenario (eulittoral), the water column scenario (pelagic) and the seafloor scenario (benthic). The field test systems have repeatably proven their stability under natural coastal marine conditions in two climate zones over many months to years of exposure. However, the 22 months of exposure time in the Mediterranean pelagic tests (Isola di Pianosa, Tyrrhenian Sea) turned out to be too short to observe strong disintegration of the exposed materials. In experiments with the same polymer types where samples were suspended from fish farm cages, [29] found much faster disintegration, presumably due to faster microbial growth at a higher nutrient content of the surrounding water.

### Samples were proven well protected against physical impact

The HYDRA^®^ test frames successfully provided protection from mechanical forces in all tests and not one sample was lost from the tests. The retrieval of almost intact sheets of the positive control material PHA after being exposed to currents and waves in the open water column for almost 2 years in the Mediterranean pelagic on the one hand, and the retrieval of heavily disintegrated, brittle samples from the SE Asia benthic impressively showed the efficiency. Previous studies used open frames without protective mesh or no frame at all to expose plastic film samples and reported complete loss of some samples during the exposure time [e.g. 19, 37]. The *gradual* material loss measured here as disintegration over time demonstrates the method chosen as suitable for long-term exposure experiments of biodegradable plastic films in the open aquatic environment.

This was also confirmed by the similar status of the test materials exposed in the mesocosm test. However, in all samples, there was a slight effect of the mesh visible at the test material surface with a differentiated thinning below the filaments and in the voids. Although the mesh was not adhering tightly there might have been some differentiations with regard to water and gas exchange, and biofilm formation. Different micro-habitat conditions could have led to the observed disintegration pattern reflecting the geometry of the mesh. However, these minor effects are seen as an acceptable trade-off for stability and operational safety of the experiments, i.e. the retrievability of the test materials after prolonged exposure time.

### Mesocosm tests can fill the methodological gap between lab and field tests

The mesocosm tests also were proven practical and generally suited to run for a longer time period in a closed circuit. The salinity could be easily adjusted, and the pH was stable over the course of the experiment. In the pelagic and benthic tests however, photosynthetic biofilms may complicate the experiments by causing a patchy overgrowth on the samples. Being of a limited amount of water and sediment, the biotic community originating from an inoculum of random diversity can lead to the dominance of few organisms (cyanobacteria and/or algae) covering large areas in each of the mesocosm tanks. In order to avoid these differences, also between tanks, we recommend conducting the experiments in the dark. Also, higher but still physiologically relevant temperatures for the respective climate zone (e.g. 25°C in the Mediterranean tests) could be used to accelerate the processes and render the tests more efficient. The heterogeneity of the disintegration between replicate samples is not to be seen as an artefact of the mesocosm tank tests but was also observed between samples in the field tests. This is interpreted as a reflection of the patchiness of the microbial community according to small-scale changes of environmental conditions, as similarly observed also for example, in marine coastal sands (e.g. [80]). This could also explain the different disintegration rates between the field experiments and mesocosm experiments with sediments from the field sites.

### Using disintegration as a proxy for biodegradation

In an open-system biodegradation test, where CO_2_ or O_2_ cannot be tracked directly with commonly available (i.e. not labelled) plastic materials, degree of disintegration is a feasible parameter to be measured in a simple and cost-efficient manner. The disintegration can however only be considered a valid proxy under the assumption that: 1) the physical forces are minimized or excluded in the open system tests and, more importantly, that the test material has shown to be biodegradable in a (standard) respirometric lab test (water: [48, 53, 54]; sediment-water interface: [49, 50]; beach sand: [51, 52, 81]). This is the case for the positive control/test material PHA Mirel P5001 that was also used in lab tests during the Open-Bio project. PHAs were also proven by others to fully biodegrade (PHB by [29] in seawater according to [48]; PHA Mirel P5001 by [82] in seawater according [48]; PHBV by [19] in seawater with biofilm from fish tank as inoculum).

As we were testing thin polymer film, we chose to measure area loss by photogrammetry instead of weight attrition [71] to assess the disintegration of the test material over time. Weight attrition might be suitable for water tests but can be easily biased by a variety of factors such as fouling organisms and adhering particles - if tests are done in contact with sediment - or by additional material loss during cleaning attempts.

By the assessment of lost area via image analysis we gained a hole/no-hole ratio after a certain exposure time. The method however intentionally ignored the third dimension of the material with the disadvantage of a technical lag phase until first holes were visible. The sometimes well discernible thinning of material was not taken into account and led to an underestimation of the disintegration at the beginning, extending the biogeochemical lag phase with a methodological lag of detectability. As soon as the material is heavily fragmented small particles might escape through the mesh leading to an overestimation of the disintegration at the very end of the experiment. Both effects will be more pronounced in the case of thicker materials. For accurate photogrammetry, standardized photos of fresh samples instead of scans of dried samples are better suited and it is recommended to put some effort into obtaining good image quality.

### Environmental relevance and the importance of a positive control: disintegration rate depends on the environmental setting

The disintegration rate of the same test material differed strongly between habitats and climate zones. A striking example is given by the comparison of the performance of the PHA samples: in the Mediterranean pelagic tests (Isola di Pianosa) there was only little material degradation after 22 months, whereas in the benthic of SE Asia (Pulau Bangka, Sulawesi) all three replicates of the same material were almost completely disintegrated after 3 months. This emphasizes the necessity of including positive controls in all tests. These data also are a valuable indication that the test results from one specific environmental setting have to be put into an ecological context and do not represent the performance of a given material in the whole marine realm. Metadata relevant for physiological activity of microbes such as temperature, oxygen, pH, light intensity, salinity, nutrients, metals, as well as granulometry, porosity and permeability of the sediments are also necessary to be able to interpret and compare disintegration data. These parameters should be measured on site and, ideally, throughout the experiment.

### Degradation under the frame differs from exposed area of interest

Due to the design of the HYDRA^®^ test frames, parts of the test material which were not intended as area of interest were covered. Only the material exposed to the matrices was analyzed as described. However, the shielding of some of the material by the frames brought along an unintentional co-testing of the material. In these conditions it can be assumed that disintegration derives solely from the action of microorganisms and that presumably only a very limited amount of oxygen is available. This gave hints for the anaerobic degradation of biodegradable plastic. Since some plastics may degrade faster under anoxic conditions, this observation should be addressed by a specific study.

### Fouling may jeopardize measurements of pelagic field tests

The specimens exposed in the pelagic test system in SE Asia were quickly covered through heavy fouling. All the materials such as frames, mesh and support structures were already overgrown after a few months by e.g. coralline algae and sponges which could not be removed without strong physical impact on the test material - a factor that had been a priority to avoid in the experimental design. Due to rather large sampling intervals, the rapid succession of the organism community and the general impact of fouling could not be studied. A small-scale observation with regular samplings every few days to weeks would be required for a good temporal resolution to understand whether fouling would hinder or accelerate polymer degradation. A previous study in the Mediterranean Sea indicated a decreased rate of disintegration by fouling [29].

### The pelagic test system is the least environmentally relevant but can deliver important insights

The habitats and locations reported in this work were chosen since they represent well the areas in which most plastic debris is found. The eulittoral test system is ideal to investigate materials with a specific density below that of seawater that initially float at the surface and can be washed ashore. This is the case for the most commonly used conventional polymers polyethylene and polypropylene, and also all items with a positive bulk buoyancy (e.g. capped bottles, foam), regardless of the material. The benthic tests represent well the seafloor scenario, the sink for most plastic materials [72, 75] due to biofilm formation. For this reason, the pelagic tests in which the materials are artificially kept from further sinking despite biofilm formation could be considered the least environmentally relevant for the study of plastic debris. However, these tests provide important information about the time during which the materials float in the water column. The pelagic test remains highly relevant to assess materials that are considered to be used directly in an aquatic environment, for example in aquaculture systems. In this case materials should be tested where they will be applied in the field.

## Conclusions

Reliable systematic tests to investigate the biodegradability of plastics in nature are urgently needed to complete the environmental risk assessment of these materials entering the marine environment. The tests presented here are an efficient toolset allowing quantitative measurements under natural aquatic conditions.

Technically, all test systems were proven to be stable and are suited for use over several years to expose and test plastic films in coastal habitats. Some maintenance to the systems such as regular cleaning is required, especially in areas of high fouling. These test systems could also easily be transferred to other aquatic habitats and test sites and be adapted to test objects instead of films. The test frames sufficiently protected the samples and no mechanical damage of the specimens was detected and we are convinced that under such test conditions the observed disintegration was caused predominantly by biodegradation. Mesocosms are well suited for manipulative experiments above lab flask scale. However, results might deviate from field experiments. Controlled parameters such as temperature and also nutrient content could be set to optimum conditions within the physiological limit of the locations the matrices are collected from. More replicates with a smaller size and the exclusion of light could lead to a more consistent data set which is less affected by heterogeneity.

Eulittoral and benthic tests are considered the most realistic scenarios for biodegradable plastics most of which have a specific density >1, and from environmental observation of the fate of conventional plastic accumulating on beaches and the seafloor. Pelagic tests are mandatory for materials which are to be used in this habitat. Testing under anoxic conditions could give a deeper insight into the performance of biodegradable polymers in aquatic environments, as most sediments are reduced in or free of oxygen.

The disintegration rate of the same test material differs between habitats and climate zones. This should be accounted for by testing in different habitats and conditions and generating a set of metadata on the ecological context of the testing site.

The outcome of this work has supported the development of a new ISO standard: “ISO 22766 (en) Plastics — Determination of the degree of disintegration of plastic materials in marine habitats under real field conditions” [47].

The assessment of marine biodegradation of plastic materials with these test systems should serve to create basic knowledge, not as an advocacy for or against a technology.

## Acknowledgements

Deep thanks go to the student assistants and interns of HYDRA for their tireless help in assembling and setting up several test systems over the years. We thank the National Park Tuscan Archipelago, Portoferraio for the access to the protected area of the Island of Pianosa with the research permit n.3063/19.05.2014. Dott. Emiliano Somigli and his staff are gratefully acknowledged for their support and for granting access to the Terme San Giovanni basin to perform the eulittoral tests. We thank Giorgio Vendetti from Hotel Mirage, Marina di Campo, for providing meteorological data (www.elbaexplorer.com). Research in Indonesia was conducted under the research permits no. 71 and 72/SIP/FRP/E5/Dit.KI/III/2017 and extensions granted by the Indonesian Government Ministry of Research, Technology and Higher Education, RISTEK-DIKTI, Jakarta to C.L. and M.W.. C.L. and M.W. express their thanks to UNSRAT welcoming them as guests researchers. Thanks to Marco Segre Reinach, Ilaria Reggi, Anna Clerici, Marco Perin and staff of Coral Eye Resort and Coral Research Outpost, Bangka Island, Sulawesi Utara, Indonesia for technical support and maintenance. Thanks also to Novamont S.p.A., Novara, Italy for providing film for tests in Italy during the Open-Bio project. Thanks to Nicolas Kalogerakis and an anonymous reviewer for valuable comments on an earlier version of the manuscript.

